# A Unified Model for Entrainment by Circadian Clocks: Dynamic Circadian Integrated Response Characteristic (dCiRC)

**DOI:** 10.1101/2021.06.16.448692

**Authors:** Zheming An, Benedetto Piccoli, Martha Merrow, Kwangwon Lee

## Abstract

Circadian rhythm is a ubiquitous phenomenon, and it is observed in all biological kingdoms. In nature, their primary characteristic or phenotype is the phase of entrainment. There are two main hypotheses related to how circadian clocks entrain, parametric and non-parametric models. The parametric model focuses on the gradual changes of the clock parameters in response to the changing ambient condition, whereas the non-parametric model focuses on the instantaneous change of the phase of the clock in response to the zeitgeber. There are ample empirical data supporting both models. However, only recently has a unifying model been proposed, the circadian integrated response characteristic (CiRC). In the current study, we developed a system of ordinary differential equations, dynamic CiRC (dCiRC), that describes parameters of circadian rhythms and predicts the phase of entrainment in zeitgeber cycles. dCiRC mathematically extracts the underlying information of velocity changes of the internal clock that reflects the parametric model and the phase shift trajectory that reflects the non-parametric model from phase data under entraining conditions. As a proof of concept, we measured clock parameters of 26 *Neurospora crassa* ecotypes in both cycling and constant conditions using dCiRC. Our data showed that the morning light shortens the period of the clock while the afternoon light lengthens it. We also found that individual ecotypes have different strategies of integrating light effects to accomplish the optimal phase of entrainment, a model feature that is consistent with our knowledge of how circadian clocks are organized and encoded. The unified model dCiRC will provide new insights into how circadian clocks function under different zeitgeber conditions. We suggest that this type of model may be useful in the advent of chronotherapies.

## Introduction

Most living organisms on the earth orchestrate their behaviors and underlying physiology in accordance with the 24-hr rotation of the earth. The cyclic environment is perceived by the organism via different cues such as the light-dark cycle, food availability, temperature changes, and social interactions. The basis for all of these daily changes is the light-dark cycle, which is therefor the predominant environmental cues (zeitgeber) are integrated by circadian clocks (Hurley et al., 2015; Young, 2018; McClung, 2019; Welkie et al., 2019; Koronowski and Sassone-Corsi, 2021). Although the key molecular players in generating the circadian rhythm signals are different, there are significant common design principles found in clocks in various model organisms (Young, 2018; Loros, 2019). Their common characteristics include an endogenous, self-sustained *circa* 24 hr (hence circadian) rhythm that persists in constant conditions, with this period length that is relative insensitive to temperature changes within a physiological range (temperature compensation) (Francois et al., 2012; Tseng et al., 2012). In entraining conditions, the clock is adjusted to the local time through synchronization. A characteristic relationship between the zeitgeber cycle and the endogenous circadian clock defines the phase of entrainment (Roenneberg et al., 2010a; Hastings et al., 2019; An et al., 2021). The robust and sustainable 24 hr period of the circadian clock, and its relative independence to temperature is intriguing, challenging the basic assumptions of biological processes. The results of mutant screens have addressed these mechanisms. We have a good appreciation of the molecular mechanisms on generating 24 hr rhythms in constant condition. In comparison, we have much less understanding of entrainment and its role in the ecology of the clock in a natural, cycling environment (Roenneberg et al., 2010a; Welkie et al., 2019).

There are two contrasting models concerning the mechanisms of entrainment, parametric and non-parametric. The non-parametric approach assumes entrainment occurs due to an instant shift of internal phase in response to the zeitgeber transition while the clock’s parameters do not reset (Comas et al., 2006). The phase shift is described by the Phase Response Curve (PRC) (Bruce and Pittendrigh, 1958; Johnson, 1992). In contrast, the parametric approach indicates that the velocity change of the internal clock caused the entrainment, expressed in a form called Velocity Response Curve (VRC) (Taylor et al., 2010). Both PRC and VRC are based on the recorded phase changes in stable conditions, usually one or more days after the light-dark transition occurs. The traditional non-parametric and parametric approaches of modeling entrainment are built on the phase-only models, meaning the only output is the phase shift.

In an attempt to integrate these two views on how circadian clocks function in a cycling environment, a model was proposed, the Circadian Integrated Response Characteristic (CiRC) (Roenneberg et al., 2010a). CiRC assumes of period matching between the endogenous clock and the zeitgeber. The organism integrates light differentially according to internal time to adjust the internal cycle length to that of the zeitgeber. Specifically, the light around dawn compresses the internal cycle, and light around dusk expands it. In comparison to the traditional approaches, CiRC has several key advantages. Firstly, CiRC makes the natural assumption of period matching. The clock actively synchronizes to the zeitgeber signal, adjusting the velocity of the internal clock. Secondly, CiRC proposes a robust way of light functioning in the entrainment process. The light stimuli accelerate or decelerate the clock according to the internal time. Thirdly, a variety of zeitgeber structures (e.g., photoperiod or thermoperiod) lend themselves to CiRC analysis and modeling. It allows us to study how the zeitgeber properties affect the entrainment. The CiRC should also reveal unique properties of various zeitgebers and their distinct signaling mechanisms.

In the current study, we develop a mathematical model reflecting the premises of both parametric and non-parametric natures of the circadian clock, dCiRC. The dCiRC faithfully describes the overt rhythms and molecular rhythms in both constant and cycling conditions. The dCiRC model describes the phase shift trajectory and the continuous velocity changes of the internal clock in a cycling condition. For the proof of concept, we analyzed 25 Neurospora ecotypes using dCiRC and found that these ecotypes have genotype specific CiRC shapes, light receptor sensitivity, and elasticity.

## Material and Methods

### Neurospora strains and cultures

The *N. crassa* ecotypes were gifts from Dr. L. Glass at University of California, Berkeley; D116, D117, D119, JW161, JW162, JW168, JW169, JW172, JW176, JW18, JW180, JW200, JW22, JW220, JW224, JW228, JW238, JW24, JW260, JW261, JW59, JW60, P4463, P4469, P4483. DBP338 (*frq*^*7*^, *bd*) was a gift from Dr. S. Crosthwaite at NIAB EMR, UK. FGSC#2489 was received from the Fungal Genetics Stock Center (Manhattan, KS). The translational FRQ:LUC reporter bearing strain X716-6a (called L3 in this report) was a gift from Dr. L. Larrondo at Pontificia Universidad Católica de Chile. The developmental rhythm was measured by Inverted Race Tube Assay as previously reported (Park and Lee, 2004). In short, we performed the experiment in two laboratory conditions, constant darkness (DD) and 12 hr light: 12 hr dark cycling condition (LD). We analyzed 25 *N. crassa* natural strains, the reference strain FGSC#2489, and DBP338 (*frq*^*7*^*;ras*^*bd*^) in DD and LD. For the chamber experiments, the inoculated race tubes are kept in a growth chamber (E-41L2, Percival) under different light conditions (DD and LD) at constant temperature 25 °C. The chambers are located in a temperature controlled (25 °C) and light-tight room. In the room, there are two different lights, white light, and red light. All race tube manipulations are performed under red light, which does not interrupt circadian regulation. Onset® HOBO® Data Loggers are used to record the temperature and light intensity in each experiment as an independent validation of the light and temperature conditions. Digital images of the race tubes were analyzed to calculate the clock properties, period and phase of entrainment, using the software Chrono (Roenneberg and Taylor, 2000) and the web-based program BioDare2 (Zielinski et al., 2014).

### Measuring molecular rhythms

Luminometry requires darkness to detect the photon emission from the cell. However, our goal is measuring the molecular clock in a light:dark cycling condition. We optimized the assay condition to measure the luciferase activity of a strain bearing a FRQ luciferase translational fusion construct in a light:dark cycling condition (S1 Fig).

### Circadian integrated response characteristic (CiRC)

CiRC describes the circadian system’s phase-dependent capacity to compress or expand its internal cycle length (denoted by *τ*_*I*_) to adjust it to that of the zeitgeber (denoted by *T*) (Roenneberg et al., 2010a; Roenneberg et al., 2010b). In terms of the velocity of the internal clock (denoted by *ω*_*I*_), CiRC represents the ability of zeitgeber signals to accelerate *ω*_*I*_ (compress *τ*_*I*_) or decelerate *ω*_*I*_ (expand *τ*_*I*_).

CiRC is composed of a sine curve and its second harmonic, see Eq (1). The second harmonic coefficient is defined as a shape factor (*s*). It determines the extent of the dead zone around noon. An asymmetry factor (*a*) determines the ratio between the amplitudes of compression (red) and expansion (blue) areas, see S2 Fig. We let *s* and *a* vary in the range [0,2] to generate CiRCs with significantly different shapes corresponding to the individuals’ various responses to the zeitgeber stimuli. The CiRC curves are normalized to an absolute maximum of 1. S2 Fig shows examples of CiRC with four combinations of *s* and *a*.

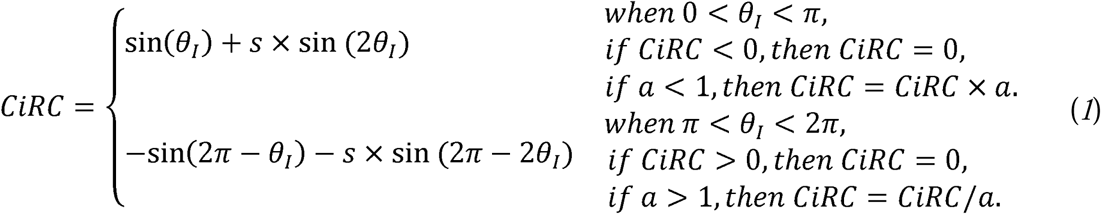

where *s* is called a shape factor, *a* is called an asymmetry factor, *θ*_*I*_ is the internal time.

### Expressing biological times: two conventions

The CiRC concept was developed using the convention ‘external time’ (ExT) and ‘internal time’ (InT) (Daan et al., 2002). The readers are encouraged to refer to the editorial paper for the full discussion on why ExT and InT were proposed. Our goal for the current study is developing ODEs that are faithfully reflecting the properties of CiRC (Roenneberg et al., 2010a). At the same time, we wanted to use ZT and CT, which are more commonly used conventions in the community. Briefly, ExT is the number of hours × 24/T elapsed since the middle of the dark period, where T is the duration of the LD cycles in hours. InT = [CT − 18]_mod24_. CT0 is aligned with ZT0 to follow the convention.

### Dynamic circadian integrated response characteristic (dCiRC) model

The dynamic CiRC model is composed of the CiRC equations and a system of ODEs to govern the behaviors of the zeitgeber cycle and the internal clock:

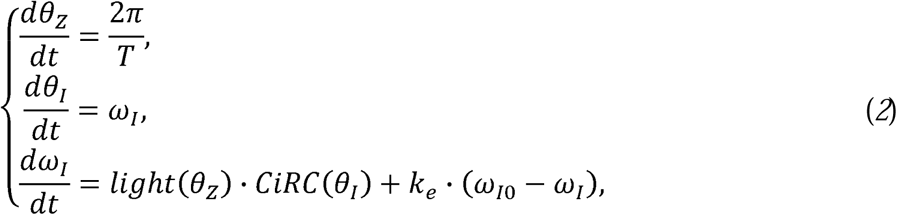

with:

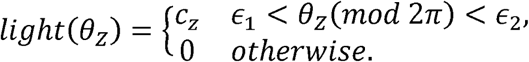

Where *θ*_*z*_ represents the phase angle of the zeitgeber cycle, *θ*_*I*_ represents the phase angle of the internal clock, *T* represents the period of the zeitgeber cycle (the time duration for the clock to complete a full cycle), *ω*_*I*_ represents the angular velocity of the internal clock, *ω*_*I*0_ represents the angular velocity of the internal clock when the organism is under constant dark (DD) condition (known as free-running), *c*_*z*_ is the zeitgeber strength which reflects the intensity of the signal and sensitivity of photo-receptors, *k*_*e*_ is an elastic constant which characterizes a frictional restoring force proportional to the velocity changes (*ω*_*I0*_ − *ω*_*I*_) of the internal clock, *ϵ*_1_ and *ϵ*_2_ indicate the time when the zeitgeber goes on and off (for example, if the organism experiences constant light, we set *ϵ*_1_ = 0, *and ϵ*_2_ = *π* which corresponds to dawn and dusk, respectively). By definition, we have that 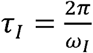 represents the period of the internal clock, and 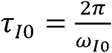 represents the free-running period (measured experimentally in DD). Although we can infer the dynamic behaviors of *ω*_*I*_ and *τ*_*I*_ from each other because they follow a reciprocal relation, we show both the changes of *ω*_*I*_ and *τ*_*I*_ because *ω*_*I*_ is a better representation for the clock running status, while *τ*_*I*_ has values around 24, which gives us a more intuitive view.

The first equation in Eq (2) governs the rate of change of *θ*_*z*_ with respect to time. The second equation in Eq (2) describes *ω*_*I*_ is the rate of change of *θ*_*I*_ with respect to time. The third equation in Eq (2) shows how the zeitgeber profile and CiRC regulate *ω*_*I*_. When there is no zeitgeber (under constant conditions), an elastic term *k*_*e*_ drives *ω*_*I*_ to converge to *ω*_*I*0_. When the zeitgeber presents around dawn (positive CiRC values), it accelerates *ω*_*I*_ (compresses *τ*_*I*_); when the zeitgeber presents around dusk (negative CiRC values), it decelerates *ω*_*I*_ (expands *τ*_*I*_).

### Phase of entrainment

Entrainment occurs when the circadian rhythm overcomes the period mismatch to adjust its period to that of the external environment (Duffy and Wright, 2005). Entrainment is characterized by a stable phase relationship with the ambient environment. The period-phase relationship between the internal clock and zeitgeber has been extensively studied in previous works (Daan and Pittendrigh, 1976; Wright et al., 2001; Duffy and Wright, 2005; An et al., 2021; Hoffmann, 2021). During the synchronization process, the phase differences (denoted by *ψ*) between the internal clock and the zeitgeber keep changing and converge to a stable value if the organism is entrained. We record the time *t*_*z*_ when 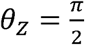, and *t*_*I*_ when 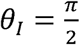(noon). Then we measure *ψ* as *t*_*z*_ − *t*_*I*_ When the organism has synchronized to the zeitgeber, the phase difference stabilizes at the value of the phase of entrainment (briefly POE, denoted by Ψ).

During the race tube experiment, a vertical marker was drawn at the growing front of mycelia at every ambient light transition, light to dark (dusk) and dark to light (dawn). The phase difference is calculated by the distance between the growing front (LD transition) and the peak of the development. We calculate the experimental POE by averaging the phase differences from the three days with minimum 3 biological replicates (Koritala et al., 2020).

### Analysis of entrainment data

#### Data processing

For developmental rhythm data: firstly, the beginning and the end of raw data are removed to exclude the outlying points. Subsequently, the truncated data is detrended using the MATLAB detrend function. We chose to subtract a quadratic trend from the data because the linear detrended results still have a clear trend that affects the following amplitude normalization process for some of the data entries. For those data entries that both the linear and quadratic detrend methods apply, we generate the final normalized data with both methods and calculate the *L*^2^ norm of the differences. The comparison shows the two methods have very similar outputs. Then, we smooth the detrended data using the MATLAB *csaps* function. In the final step, we fit the peak heights of the smoothed data with polynomial equations and normalize the amplitude of the data to 1. For molecular rhythm data, we follow the same process, with a slight modification. We keep all data points and apply the cubic detrend method to all data entries.

### Determination of dCiRC parameters

To determine the best-fitting dCiRC parameters for each genotype, we fit the simulated dCiRC phase trajectory to the processed data and solve a convex optimization problem. We apply the MATLAB PSO (Particle Swarm Optimization) algorithm with the following convex cost function:

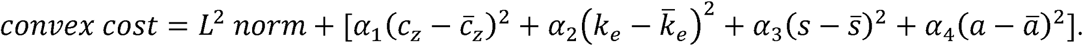

Where *c*_*z*_ ∈ [0,3] is the zeitbeger strength, *k*_*e*_ ∈ [0,3] is the elastic factor and the unit of *k*_*e*_ is *s*^−1^, *s* ∈ [0,2] is the shape factor, *a* ∈ [0,2] is the asymmetry factor, the *L*^2^ *norm* is the square root of the sum of the square differences between the simulated and experimental data points, 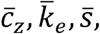 and *ā* are reasonable targeting values (estimated from data) for *c*_*z*_, *k*_*e*_, *s*, and *a, α*_1_, *α*_2_, *α*_3_, *α*_4_ are coefficients of the convex terms. We choose a reasonable range for the parameters based on the testing simulations.

For all genotypes, we fit the data and take the average 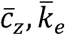 of all fitting results as the targeting values. Then, we fit all three data entries of each genotype. We calculate the best-fitting *c*_*z*_, *k*_*e*_ values by averaging three pairs of *c*_*z*_, *k*_*e*_ values. Then we explore the 3D space of “convex cost – s – a” and find the pair of (*s,a*) values that minimizes the convex cost. The pair of (*s,a*) that minimizes the total cost for all three data entries is the best fitting *s* and *a* for this genotype. Then we plot the best fitting CiRC and the change of ω_I_ over time for each genotype. We also simulate Ψ and compare it to the one observed from experiments. Table 1 shows the key parameters we used in the fitting process.

**Table 1:**
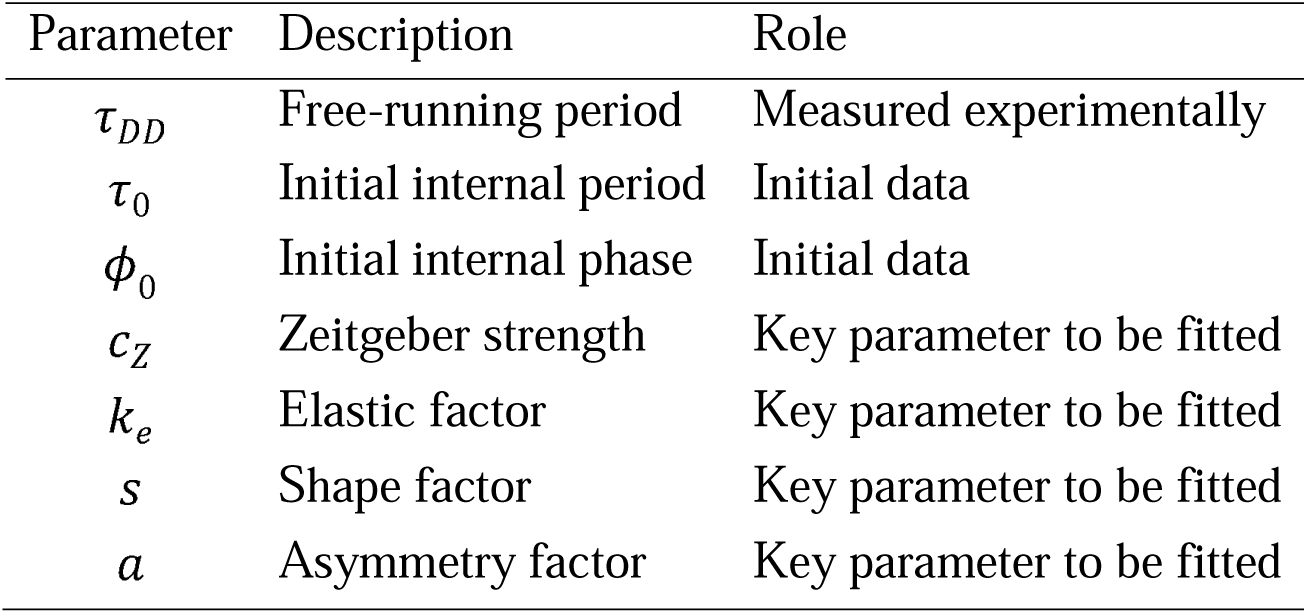
Biological meanings and the roles of the main parameters in the dCiRC model.

| Parameter | Description | Role |
| --- | --- | --- |
| $\tau_{DD}$ | Free-running period | Measured experimentally |
| $\tau_0$ | Initial internal period | Initial data |
| $\phi_0$ | Initial internal phase | Initial data |
| $c_z$ | Zeitgeber strength | Key parameter to be fitted |
| $k_e$ | Elastic factor | Key parameter to be fitted |
| $s$ | Shape factor | Key parameter to be fitted |
| $a$ | Asymmetry factor | Key parameter to be fitted |

## Results

### The velocity of the clock (*ω*_*I*_) and phase of entrainment (Φ) in dCiRC

To determine if dCiRC (Materials and Methods) reflects both the change of the circadian clock velocity and the phase of entrainment, we simulated the dCiRC model under different light conditions (Fig 1). The light profile was composed of two rounds of zeitgeber cycles which were repeated twice. In each of the 35-day rounds, the first 20 days were a 24h zeitgeber cycle (12h:12h, light on at ZT0, light off at ZT12) conditions, and the remaining 15 days were under DD conditions. We choose the CIRC variant in Fig. 1A with *s* = 0, *a* = 1. The rest of the parameters were: *τ*_*I*0_ =23*h, T=* 24*h, c*_*z*_ = 0.025,*k*_*e*_ =0.05. There were four stages in each round of zeitgeber cycles. In the beginning, the clock had an initial *τ*_*I*_ = *τ*_*I*0_ = 23*h*. From day 1 to day 11 is the “synchronizing transition” stage (briefly ST), the organism goes through the “synchronizing” process. During each day, the light signal in the early day (ZT0 – ZT6) compresses *τ*_*I*_ (accelerates *ω*_*I*_), the light signal in the late day (ZT6 – ZT12) expands *τ*_*I*_ (decelerates *ω*_*I*_). The phase difference converges to Ψ. From day 12 to day 20 is the “synchronizing stable” stage (SS), the organism is “Synchronized” under cycling conditions. *τ*_*I*_ stays in a dynamic equilibrium around *T* = 24*h*, and the phase difference stabilizes at Ψ = 2.96*h*. The following is the “desynchronizing transition” stage (DT), the organism is desynchronizing from the zeitgeber clock during day 21 through 27 because it is in DD. Here, the phase differences keep changing on each simulated day only due to the period mismatch. From day 28 to day 35 is the “desynchronizing stable” stage (DS), the organism is in free-running status where the phase differences stabilized and *τ*_*I*_ stays at the value of *τ*_*I*0_ =23*h*. The phase difference continues changing due to the period mismatch. From day 36 to day 70, the same procedure is repeated. The organism experiences the same stages while the trajectories of *τ*_*I*_, *ω*_*I*_, and *ψ* have shifted.

**Figure 1.**
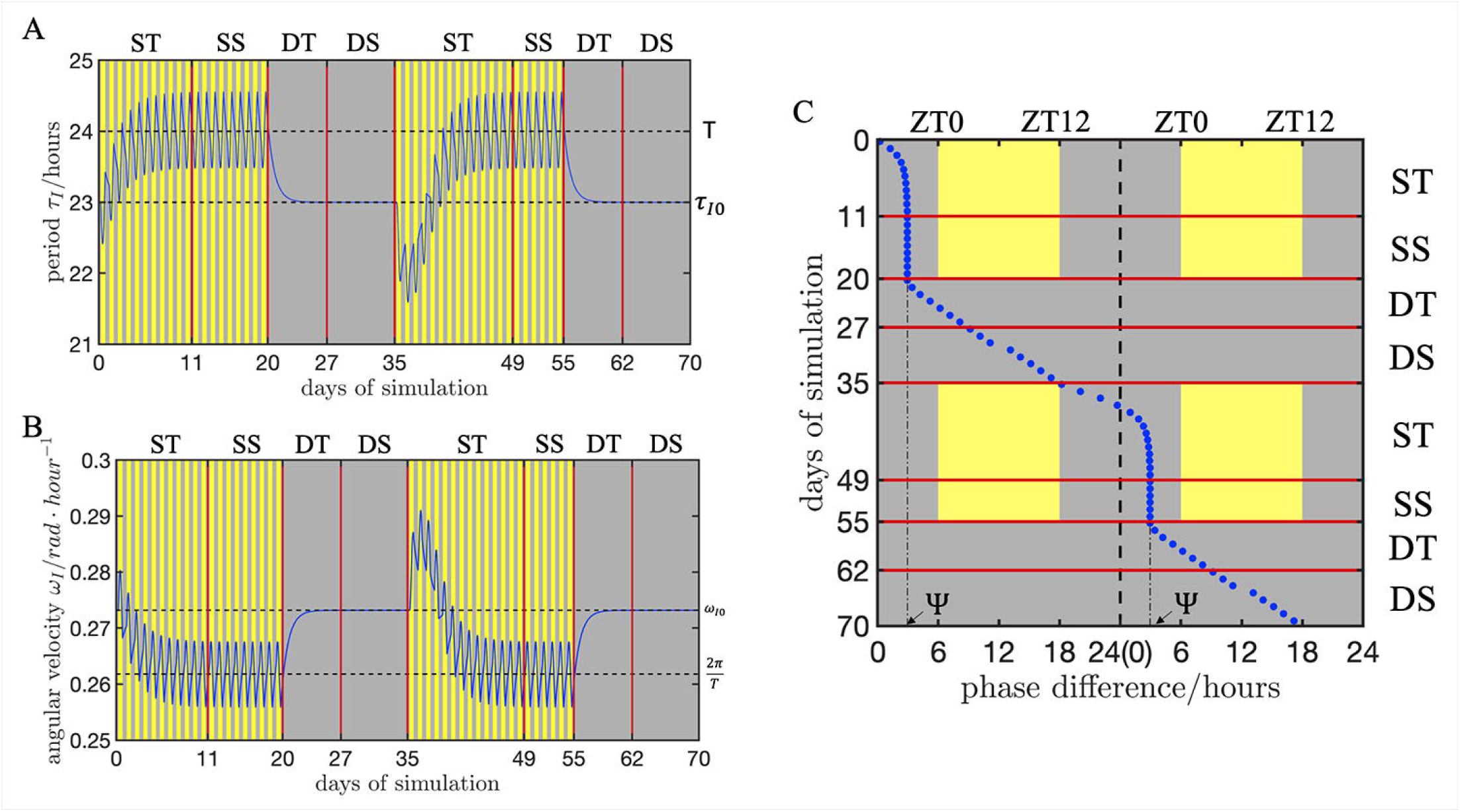
Simulations of the dCiRC model showing the clock’s velocity (*ω*_*I*_) and the phase of entrainment (Ψ). An illustrative example of the dCiRC model reflecting both the parametric and non-parametric aspects of the entrainment. In the experiment, the light profile was composed of two rounds of zeitgeber cycles which were repeated twice. In each of the 35-day rounds, the first 20 days were a 24h zeitgeber cycle (12h:12h, light on at ZT0, light off at ZT12) conditions, and remaining 15 days were under DD conditions. There are four stages in each round of zeitgeber cycles: synchronizing transition (ST), synchronizing stable (SS), desynchronizing transition (DT), and desynchronizing stable (DS). (A) The trajectory of the internal clock’s period (*τ*_*I*_) change. T and *τ*_*I*0_ represent the zeitgeber period and the free-running period, respectively. (B) The trajectory of the internal clock’s velocity (*ω*_*I*_) change. 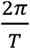 and *ω*_*I*0_ represent the velocity of the zeitgeber clock and the free-running velocity of the internal clock, respectively. (C) Double plot showing the trajectory of the phase differences (*ψ*) between the internal clock and the zeitgeber clock. We choose the CiRC type in Figure 1A with *s* = 0, *a* = 1. The rest of the clock’s parameters are: *τ*_*I*0_ = 23*h,T* = 24*h, c*_*z*_ = 0.025, *k*_*e*_ = 0.05.

### Genotype-specific dCiRC parameters

Next, we aimed to identify the clock parameters based on the experimental data in entraining conditions. For this, we assayed the overt daily rhythm of asexual development in 25 *N. crassa* natural ecotypes and a long-period mutant DBP338 (*frq*^*7*^*;ras*^*bd*^, *τ*_*I*0_ = 30.0*h*) under DD and LD conditions (Materials and Methods, S1 Data). We used these experimental data to find the best fitting parameters for the dCiRC model (Fig 2). Due to genetic variations, complex environmental cues, limits of recording techniques, and artifacts, the recorded rhythmic data contains many different shapes. On the other hand, the data diversity among all genotypes also indicates the natural variations of initial phase at the beginning of the experiment, the non-uniform light responses, and so on. We truncate the parts of data that do not follow the consistent rhythmic behaviors which cause different time spans of fitting curves. We also fitted the initial phase for each genotype which cause the shift of the fitting curves.

**Figure 2.**
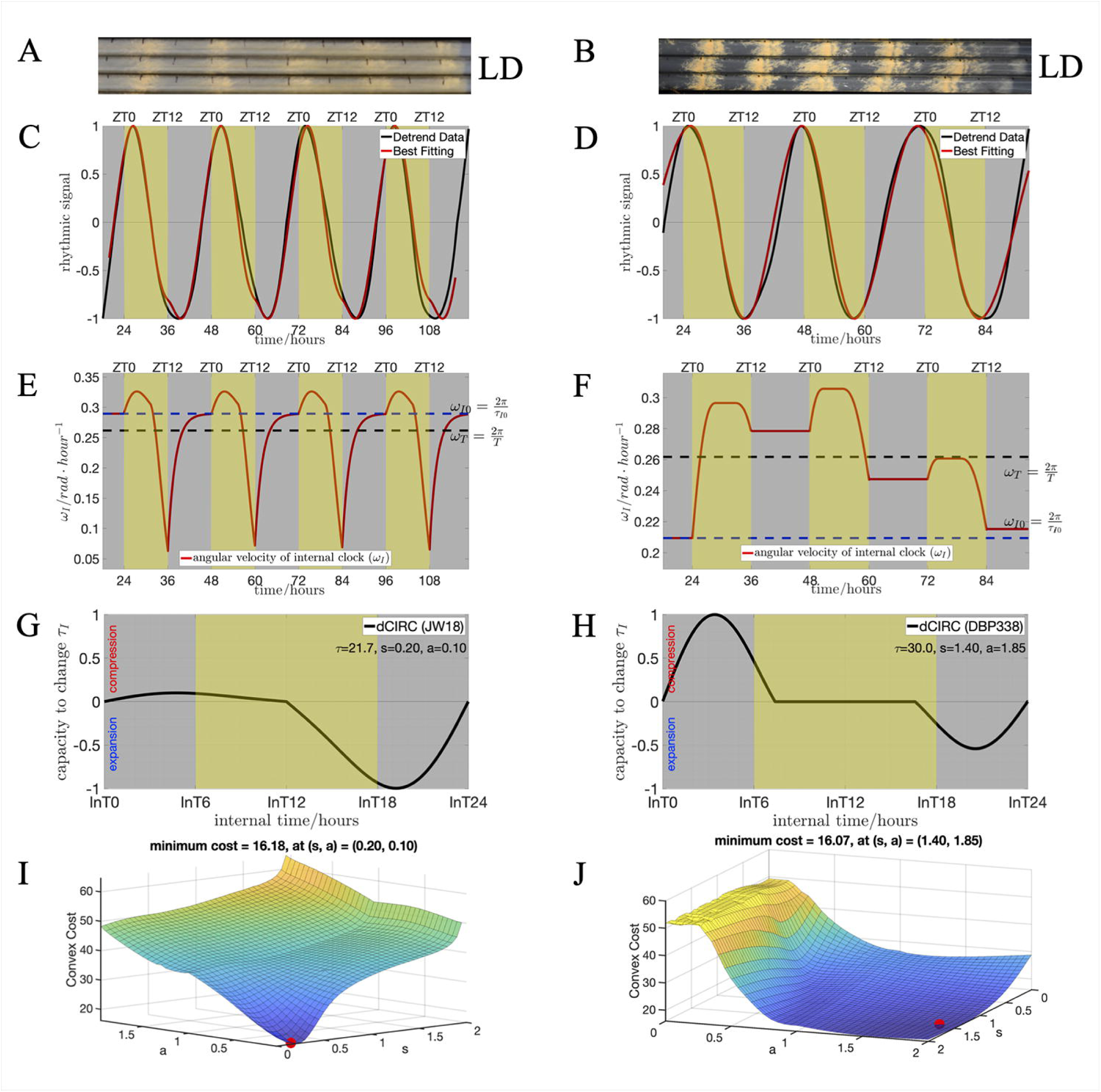
Using developmental rhythms data of *N. crassa* to find the genotype-specific dCiRC parameters. The organisms are exposed to light from ZT0 to ZT12 on each day. (A) and (B) are the race tube images showing the conidiation process of ecotypes JW18 (with *τ*_*I*0_ = 21.7*h* < 24*h*, short period), and DBP338 (with *τ*_*I*0_ = 30.0*h* > 24*h*, long period) under LD laboratory condition, respectively. (C) and (D) are the results of using dCiRC model to fit the phase trajectory for ecotypes JW18 and DBP338, respectively. The black curves represent the processed rhythmic data of conidiation (detrending, smoothing, and normalization). The red curves are best fitting curves find by the dCIRC model with convex optimization algorithms. The data is shown in both the experimental time (bottom axis) and zeitgeber time (ZT, top axis). The best-fitting clock’s parameters for JW18 are: *c*_*z*_ = 0.2658, *k*_*e*_ = 0.4019, *s =* 0.20, *a =* 0.10. The best-fitting clock’s parameters for DBP338 are *c*_*z*_ = 0.0394, *k*_*e*_ = 0.0056, *s =* 1.40, *a =* 1.85. (E) and (F) show the changes of the clock’s velocity during the entrainment process for ecotypes JW18 and DBP338, respectively. 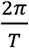 and *ω*_*I*0_ represent the velocity of the zeitgeber clock and the free-running velocity of the internal clock, respectively. (G) and (H) are the genotype-specific dCiRC curves for ecotypes JW18, and DBP338, respectively. (I) and (J) are results of using the 3-D convex cost space generated by the dCiRC model to identify the best-fitting combinations of (*s, a*) for ecotypes JW18, and DBP338, respectively. The blue color corresponds to the smaller costs, and the green/yellow color corresponds to the larger costs. The red dot represents the best-fitting (*s, a*) values that minimized the convex cost.

In conventional race tube analysis, the phase data is calculated by averaging the phase from each day during the experiments (Materials and Methods). The averaged phase value does not reflect the status of the entrainment process. We observed that the peaks of the detrended data of JW18 present roughly the same ZT hour at each day, whereas those of the detrended data of DBP338 keep advancing (Fig 2C and D). This suggests both that ecotype JW18 reaches a stable entrainment in four days of experimental data, whereas the long period strain DBP338 does not. The light signal during ZT0 - ZT6 is used by the internal clock to overcome the period mismatch (between the *θ*_*I*_ and *θ*_*z*_) and the effect of *k*_*e*_ to increase *ω*_*I*_ (Fig 2E). The light signal present between ZT6 - ZT12 is integrated with the effect of *k*_*e*_ to decelerate *ω*_*I*_. During the ZT12 - ZT24 period, the elastic effect drives *ω*_*I*_ to converge to the free-running velocity. The stationary point of the velocity is by definition at noon. The periodic behaviors of the internal clock’s velocity indicate that the strain is stably entrained with *c*_*z*_ = 0.2658 *and k*_*e*_ =0.4019. The internal clock’s velocity of DBP338 changes with the same mechanisms while no periodic behaviors are observed (Fig 2F). Instead, the internal clock of DBP338 runs at different rates at the end of each day. This suggests that the clock is not yet entrained with *c*_*z*_ = 0.0394 *and k*_*e*_ = 0.0056. The flat zone around ZT12 is cause by the extensive dead zone in the dCiRC (see Fig 2F). A small value of the elastic factor allows the clock to run at nearly stable rates during ZT12 - ZT24. We concluded that the larger *c*_*z*_ *and k*_*e*_ values reflect the DBP338’s insensitivity to the light. Thus, the internal clock’s velocity fluctuates in a narrower range in comparison to the strain with a shorter *τ*_*I*0_. As expected, each genotype has a specific dCiRC curves (Fig.2G and 2H) with unique values of s and a (Fig 2I and J, best-fitting parameters of all 26 genotypes in S1 Data). For genotypes with short *τ*_*I*_, 14 out of 25 (56%) genotypes with short *τ*_*I*_ have a combination of *c*_*z*_ < 0.6,*k*_*e*_ > 0.6 (S1 Data). The results indicate that there exists a relatively strong restoring force to keep a self-sustained free-running oscillation, while a relatively weak entrainment strength of *c*_*z*_ is shown. The best-fitting (*s,a*) combinations show large variations among all genotypes. The *s* values distribute across the range of [0,1.4], while 23 out of 25 (92%) genotypes have *a* values less than 0.40 (S1 Data).

We calculated the areas under the dCiRC curve with a sign (the area is positive from 0h to 12h, and negative from 12h to 24h) and defined it as the total area under curve (TAUC) of dCiRC (S3 Fig). We define the portion of the above signed area covered by light signal as the light-exposed area under curve (LAUC) (S3 Fig). Positive values of TAUC and LAUC reflect that the compression area is larger than the expansion area, thus the overall light effect is compressing *τ*_*I*_. Negative values of TAUC and LAUC reflect that the expansion area is larger than the compression area, thus the overall light effect is expanding *τ*_*I*_. For JW18, the optimized combination of (*s,a*) is (0.20,0.10). The expansion area being larger than the compression area (negative TAUC and LAUC) is consistent with the expectation for the genotype with shorter period (*τ*_*I*_ = 21.7*h* < 24*h*). For DBP338, the optimized combination of (*s,a*) is (1.40,1.85). There is an extensive flat zone around noon. This implies that *ω*_*I*_ actively changes in response to the light signal around dawn or dusk while the zeitgeber around noon is ineffective. Its dCiRC has larger compression area than expansion area (positive TAUC and LAUC) which is consistent with our expectation for the genotype with longer period (*τ*_*I*_ = 30.0*h* > 24*h*). The *a* value represents the compression/expansion ratio. When *a* < 0.20, the expansion effect is more than five times that of the compression effect (negative TAUC and LAUC). For genotype DBP338 with *τ*_*I*_ > 24*h*, a large *a* value of 1.85 indicates that the compression effect is almost the double of the expansion effect (positive TAUC and LAUC). However, it has relatively small *c*_*z*_ = 0.0394 and *k*_*e*_ = 0.0056. This implies that the DBP338 strain shows less sensitivity and resistance when responding to the light signal. All fitting results meet our expectation that *ω*_*I*_ decelerates in response to the zeitgeber when an organism has a short *τ*_*I*_, and *ω*_*I*_ accelerates when an organism has a long *τ*_*I*_.

### Analyzing molecular rhythm data

We optimized the luciferase reporter assay for N. *crassa* under a cycling environment (Materials and Methods, S1 Fig). Unlike the developmental rhythm, the luciferase reporter data reflecting the molecular rhythm are regular and consistent (Fig 3). In the beginning of the entrainment process, the internal clock shows a small trend of delayed phase; small absolute values of slope of the tangent line (Fig 3A). This observation agrees with the prediction that the internal clock starts with velocity 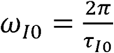 and encounters a gentle decrease (from ZT0 to ZT2) (Fig 2B). In each successive day of the light exposure, the morning light increases *ω*_*I*_ and the afternoon light decreases *ω*_*I*_. This resembles in some sense an iterative process. Starting from the fourth day of light exposure, we observed a significant change of *ω*_*I*_ from ZT0 to ZT2. This change suggests the light has shifted the trend of deceleration to acceleration. Stable entrainment is indicated when *ω*_*I*_ is synchronized to reach *ω*_*T*_ for the first time. This implies that the organism is not completely entrained but will be soon. The dCiRC of the molecular rhythm shows a larger compression area under light exposure (Fig 3C). The larger compression area than the expansion area (negative TAUC and LAUC) is consistent with the expectation for the genotype with longer period (*τ*_*I*0_ = 24.7*h*). There is no flat zone around noon, which tells us that the organism makes full use of the light signal in order to be entrained.

**Figure 3.**
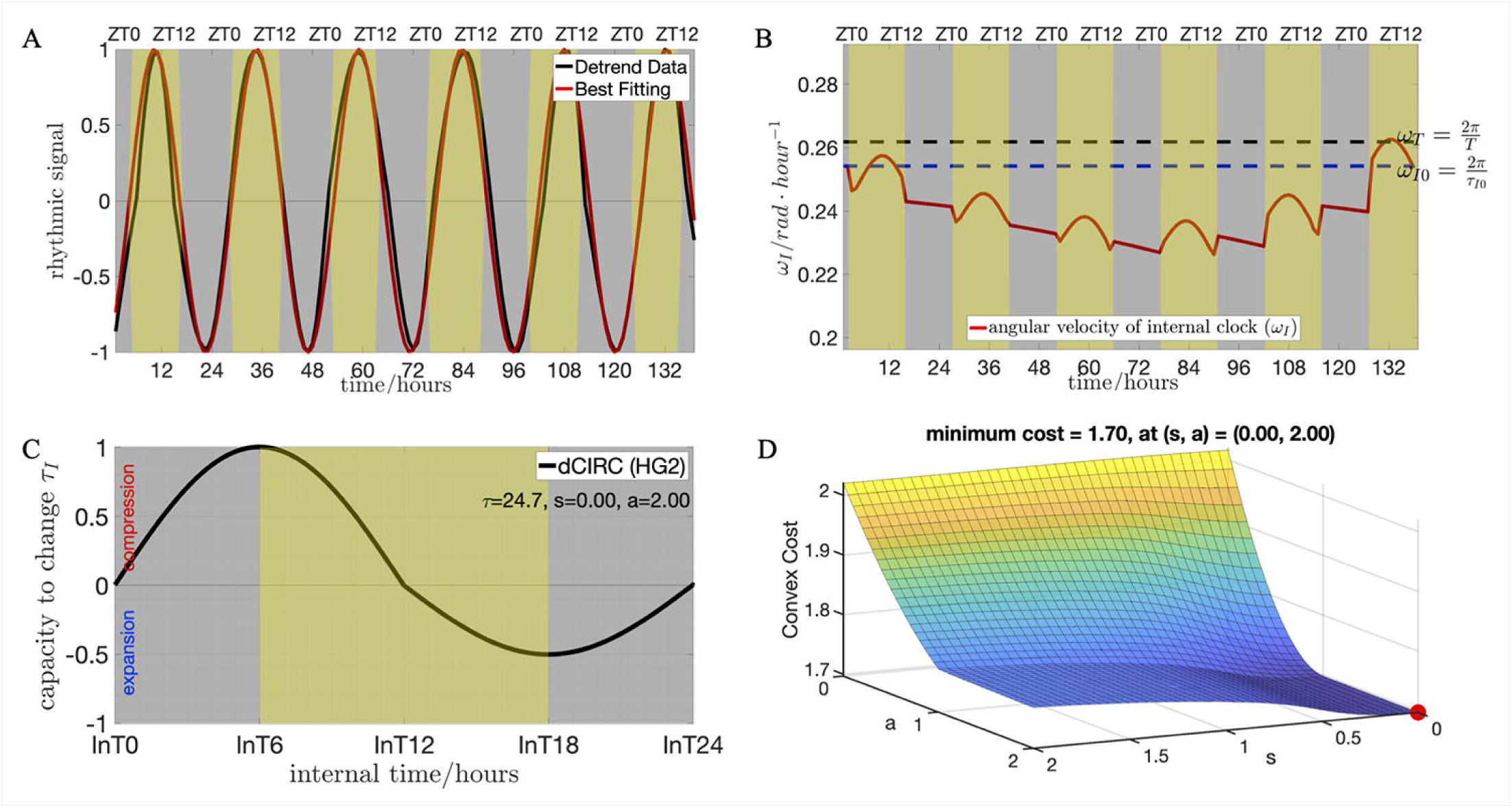
Examples of using the dCiRC model to fit the molecular rhythm data. The data recorded the luciferase activity of FRQ in *N. crassa*. The organism (*τ*_*I*0_ = 24.7*h*, long period) is exposed to light from ZT3 to ZT15 on each day. (A) The results of using the dCiRC model to fit the phase trajectory data. The black curve represents the processed rhythmic data. The red curve is the best fitting curves find by the dCiRC model with convex optimization algorithms. The data is shown in both the experimental time (bottom axis) and zeitgeber time (ZT, top axis). The best-fitting clock’s parameters are *c*_*z*_ = 0.0287, *k*_*e*_ = 2.7933, *s =* 0.00, *a* = 2.00. (B) The changes of the clock’s velocity during the entrainment process. 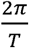 and *ω*_*I*0_ represent the velocity of the zeitgeber clock and the free-running velocity of the internal clock, respectively. (C) The genotype-specific dCiRC curve. (D) The 3-D convex cost space generated by the dCIRC model to identify the best-fitting combinations of (*s, a*). The blue color corresponds to the smaller costs, and the green/yellow color corresponds to the larger costs. The red dot represents the best-fitting (*s, a*) values that minimized the convex cost.

### Results of predicting phase of entrainment by dCiRC

The non-parametric property of the dCiRC model allows us to predict the phase of entrainment. For the purpose of measuring the accuracy of the prediction, we compared the simulated phase of entrainment (Ψ_*sim*_) with the experimentally measured phase of entrainment (Ψ_*exp*_), then we calculate the model’s predictive power (Fig.4). In this experiment, we used the software Chrono to calculate Ψ_*exp*_ by taking the average *ψ*_*exp*_ values (Materials and Methods). We calculate Ψ_*sim*_ in the same way by taking the average *ψ*_*sim*_ values. We found that dCiRC provides a high accuracy of prediction; the linear regression line of Ψ_*sim*_ (Ψ_*exp*_) with a slope of 0.2467 (0.1984) (Fig 4A). We define the predicting errors (*ϵ*) by subtracting Ψ_*exp*_ from Ψ_*sim*_. The small values of absolute mean of *ϵ* and C.I. indicate that the dCiRC predicted POEs fit the experimental data well (Fig 4B). We expected an accurate prediction result because we used the PSO algorithm and the L2 norm cost function. The dCiRC model is a reliable tool for processing the biological rhythmic data and predicting and calibrating the POE results. We also define the internal clock with large zeitgeber strength (beyond 75th percentile) as the weak oscillator, the internal clock with small zeitgeber strength (below 25th percentile) as the strong oscillator (S4 Fig). Our analysis show that the organisms with weak oscillator results in a more extensive range of POE (Ψ_*sim*_) compare to the organisms with strong oscillator (Granada et al., 2013).

**Figure 4.**
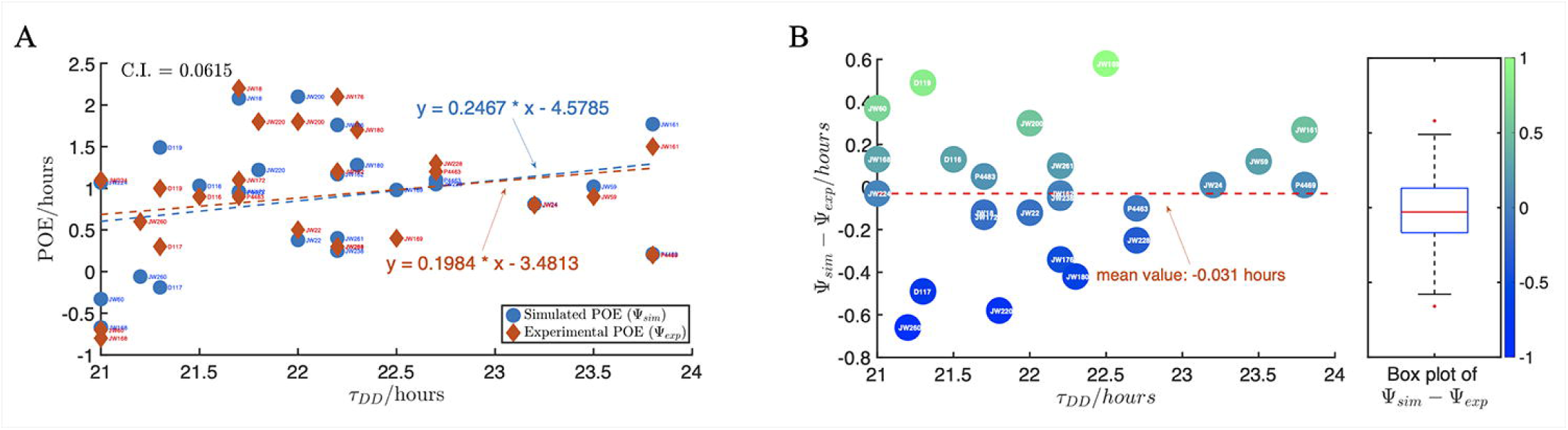
Predicting phase of entrainment by the dCiRC model. (A) The comparison of the simulated POE (Ψ_*sim*_) and the experimental POE (Ψ_*exp*_) showing the high prediction power of the dCiRC model. The round blue markers represent Ψ_*sim*_ values predicted by the dCiRC model, and the red diamond markers represent Ψ_*exp*_ values obtained from the experiment. The blue (red) dotted line is the linear regression line of Ψ_*sim*_ (Ψ_*exp*_) with a slope of 0.2467 (0.1984). We define the confidence level (C.I.) as the L-2 norm of *ϵ* = Ψ_*sim*_ − Ψ_*exp*_ divided by the number of ecotypes, 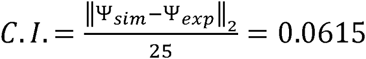. (B) Distribution of POE predictability by dCIRC of ecotypes based on the *τ* _*DD*_. The left side is a scatter plot of *ϵ* and *τ*_*DD*_, the right side is a boxplot to visualize the summary statistics of *ϵ* for all 25 short *τ*_*I*_ ecotypes. The color bar represents the absolute values of the *ϵ*. The absolute mean of *ϵ* is 0.031*h* ≈ 1.86*min*.

### Non-uniform response of ecotypes in cycling condition

We hypothesized that the parameters in dCiRC of the ecotypes that we have characterized in the current study reflect an evolved adaptation of the circadian clock to its local habitat. Strains are collected from geographically diverse locations. To test this hypothesis, we firstly analyze the relationships between TAUC and LAUC. LAUC value reflects how the ecotype responds to the current light exposure while TAUC reflects the ability of the ecotype responds to all potential light exposures (Material and Methods). There exists a strong positive linear relationship between TAUC and LAUC (r = 0.91) such that we can use LAUC to represent both the current and potential light responses in the following analysis. This is also important when we predict how one ecotype responds to various light conditions (Fig 5A).

**Figure 5.**
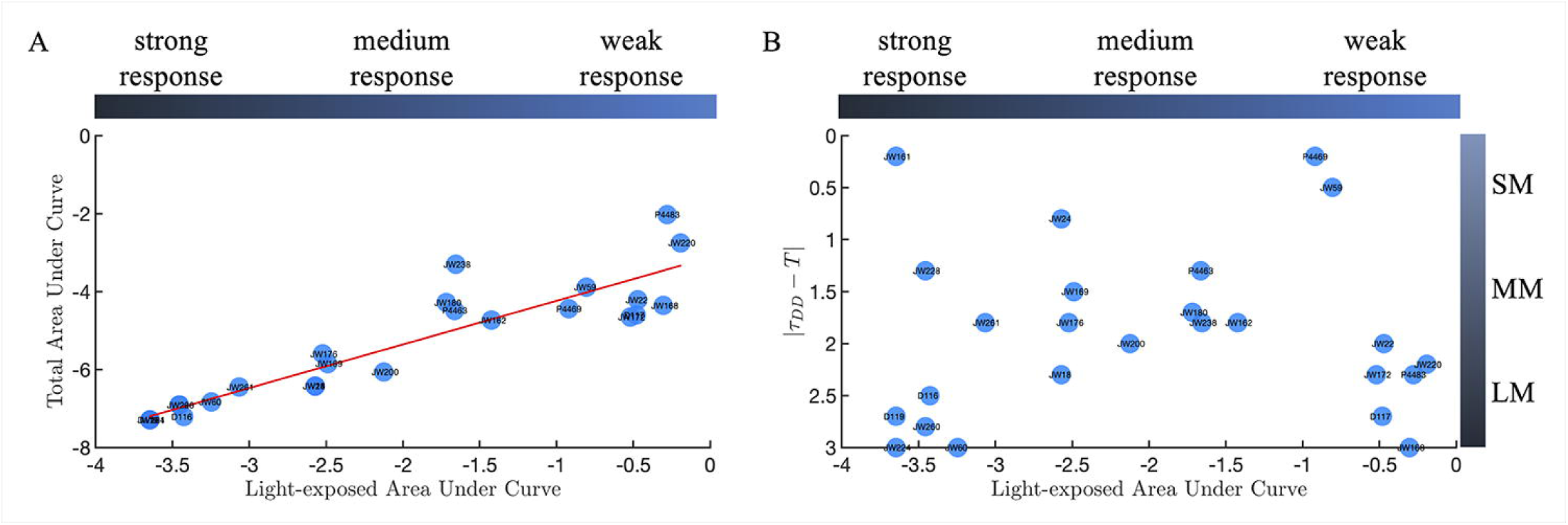
Non-uniform response of ecotypes in cycling condition. (A) Scatter plot of TAUC and LAUC with the linear regression line (in red). The color bar (top) represents the strength of light response according to the absolute LAUC values. The light response ability is categorized in the following: weak response when 0 ≤ |*LAUC*| ≤ 1, medium response when 1 < |*LAUC*| ≤ 3, strong response when 3 < |*LAUC*| ≤ 4. (B) Scatter plot of LAUC and |*τ*_*DD*_ − *T*| showing how the strength of light response is related to the period mismatch. The horizontal color bar (top) represents the strength of light response according to the absolute LAUC values. The vertical color bar (right) represents the period mismatch (|*τ*_*DD*_ − *T*|). The period mismatch is categorized in the following: small period mismatch when 0 ≤ |*τ*_*DD*_ − *T*| ≤ 1, medium period mismatch when 1 < |*τ*_*DD*_ − *T*| ≤ 2, strong period mismatch when 2 < |*τ*_*DD*_ − *T*| ≤ 3.

We define the light response ability for each genotype according to its LAUC value. The sign of LAUC indicates whether the overall light effects is compressing or expanding the internal period (*τ*) (Material and Methods). The absolute value of LAUC reflects the strength of the light response. In Fig.5A, we define that when 0 ≤ |*LAUC*| ≤ 1, the ecotypes have weak response to the light (8 of 25 ecotypes); When 1 < |*LAUC*| ≤ 3, the ecotypes have moderate response to the light (9 of 25 ecotypes); When 3 < |*LAUC*| ≤ 4, the ecotypes have strong response to the light (8 of 25 ecotypes). The results show that all ecotypes evenly distribute among the three light response groups. To study the relationship of light response and period mismatch, we calculate the value of |*τ*_*DD*_ − *T*| for each ecotype and define three groups. When 0 ≤ |*τ*_*DD*_ − *T*| ≤ 1, the ecotypes have small period mismatch (SM) to the zeitgeber; When 1 ≤ |*τ*_*DD*_ − *T*| ≤ 2, the ecotypes have medium period mismatch (MM) to the zeitgeber; When 2 ≤ |*τ*_*DD*_ − *T*| ≤ 3, the ecotypes have large period mismatch (LM) to the zeitgeber. In fig.5B, we observe that for ecotypes with strong and weak light responses, their |*τ*_*DD*_ − *T*| values are widely distributed in the range of 0h to 3h. The ecotypes with medium light response have a narrow range of |*τ*_*DD*_ − *T*|. Interestingly, we also find that in each period mismatch group (SM, MM, and LM), the ecotypes utilize all three different light response strategies. The results suggest that there is no significant relationship between light response and period mismatch.

### dCiRC model reproduces PRCs

PRC has been the main means of appreciating the phase shift by a certain zeitgeber of a circadian clock under a constant condition. We predicted that the dCiRC model should be able to produce PRC using the best-fitting parameters in dCiRC (S1 Data). In addition, we can use the dCiRC model to study the phase response curves for various duration, timing, and intensity of the light. More importantly, the reverse of the above process allows us to utilize the massive data of PRCs to generate the dCiRC curves. This gives us potential to build a “circadian fingerprint” database composed of genotype-specific or species-specific dCiRC parameters that reflect the entrainment mechanisms for the organisms.

As an example, we used the dCiRC model to generate the phase response curve to one hour of light exposure. Remarkably, that the dCiRC-generated PRC reflects the key features of the traditional experimental-generated PRC: a small portion of the advanced zone in the dawn is followed with a dead zone around the noon. In the afternoon, a delayed zone, and a transition from the phase delays to advances is represented. A day ends in a zone of phase advances. For each genotype, the PRC shows similarities to its specific dCiRC. For genotype JW59 with *τ*_*I*_ = 23.5, a delayed phase of the internal clock is required in order to entrain. As expected, the PRC shows that the amplitude of the delayed zone is larger than that of the advanced zone as we expected. In contrast, for genotype DBP338 with *τ*_*I*_ = 30.0*h* the PRC shows a larger advanced zone relative to delays to reach an advanced phase shift (Fig 6).

**Figure 6.**
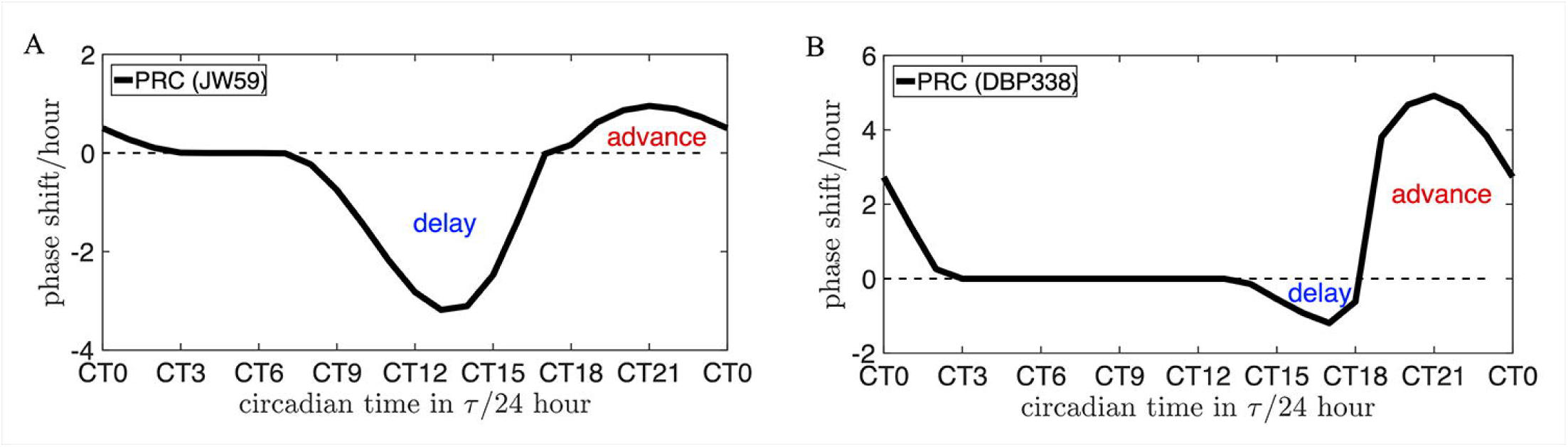
Examples of using the dCiRC model to generate PRCs. (A) The phase response curve of ecotype JW59 to one hour of light exposure. (B) The phase response curve of ecotype DBP338 to one hour of light exposure. The horizontal axis is the time of the light exposure measured in circadian hours (CT).

## Discussion

There is a large number of studies focused on modeling the circadian clock and predicting entrainment, and they are categorized into parametric and non-parametric approaches. We have generated an ODE-based model that reflects the characteristics of both parametric and non-parametric understanding of the circadian clock. One of the dCiRC model’s main features is that the circadian system inherits a phase-dependent capacity to compress or expand its internal period to match the period of the zeitgeber cycle. This feature reflects the parametric approach and allows the dCIRC model to describe the underlying changes in clock parameters. The other important feature of dCiRC is predicting phase shift trajectory during entrainment. This is the non-parametric aspect of the dCiRC model which allows us to study the period-phase relationship.

Velocity Response Curve (VRC) is designed to capture the phase shift features of a PRC and the VRC has similar shape relative to a PRC. Taylor et al. proposed a phase-only model based on the modified evolution of the internal phase depending on the light signal and the VRC (Taylor et al., 2010). While the idea has some similarities with the dCiRC approach, there is a substantial difference at the mechanistic scale: in the VRC model, the phase adjustment is described by a single equation; thus, light instantaneously changes the internal clock’s angular velocity. In contrast, in the dCiRC model, light affects the internal clock’s angular acceleration. Theoretically, the high complexity allows the dCiRC to capture the transient dynamics for changes from LD to DD. We elaborate the statement in the following two aspects. Firstly, the “high complexity” is partially reflected by the inclusion of four parameters *c*_*z*_, *k*_*e*_, *s*, and *a*. It allows us to describe the velocity changes in the presence of light (acceleration and deceleration) as well as the relaxation (recovery) process in constant dark. While the VRC model has instant recovery, the advantage of using the dCiRC model to study after-effects supports our statement of “capture the transient dynamics”. Secondly, let’s recall a few remarks from another VRC paper. The original VRC idea was introduced as a theoretical construct to describe how the amount of “advance or speed-up or retardation or slow-down is a function of the area under the VRC curve and the light intensity”(Swade, 1969). This idea incorporated the phenomenon that “animals somehow average or sum the light intensities they see over a period of time each day, and that this average or total is the entraining agent” (Swade, 1969). In the dCiRC model, the light and CiRC curve will change the angular velocity (*ω*_*I*_) via the integration of 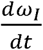 during a small increment of time, which is conceptually consistent with the idea that the “entraining agent” takes effects upon a duration of time.

Furthermore, we compare the two models from the perspective of predicting entrainment. In Taylor et al., the authors estimated a VRC from an existing PRC and subjected it to the entrainment data under cycling conditions. As we know that PRC recorded the phase shifts in DD after the light exposure, the VRC approach has an inherent deficiency of utilizing information from DD condition (PRC) to study entrainment in LD conditions. In contrast, the dCiRC approach estimated a dCiRC directly from the entrainment data in LD conditions and simultaneously studied how the organism adjusts its internal clock’s parameters based on the dCiRC in response to the light signal. This suggests that our approach in dCiRC better captures the parametric nature of the clock (Taylor et al., 2010).

Compared to the existing mathematical modeling of the circadian clocks, a major advantage of the dCiRC model is the application to natural cycling conditions. For instance, Abraham 2010 et. al. proposed a Poincaré oscillator-based model to study how the phase of entrainment is related to the mismatch between the internal and zeitgeber period (Abraham et al., 2010). A two-step model in a previous study (An et al., 2021) and the dCiRC model in the current study (S1 Data) suggest that there exists a complex relationship between period and phase rather than a positive linear relationship by fitting phase data under LD conditions. Furthermore, due to the unique properties of our unified theory, the dCiRC model can extract information from the non-parametric aspects (such as phase of entrainment) and reveal the mechanisms of underlying parametric aspects (such as velocity) of the circadian systems. (Granada et al., 2013) concluded that a weak oscillator (with large zeitgeber strength) results in a more extensive range of POE compared to a strong oscillator (with small zeitgeber strength). Our data support this conclusion (S1 Data and S4 Fig). In addition, the dCiRC model illustrates the evolution of the internal clock’s velocity changes. The results gave us insights into how the organisms with similar internal periods have significantly different responses to the light, and as a consequence, result in various phases of entrainment.

We observe that there is no significant relationship between the free-running period and light response (Fig 5). One explanation of diverse light response behaviors is that the initial status of the internal clock plays a vital role in the entrainment process. This suggests that organisms have a complex system to determine their entrainment strategy. The dCiRC model provides an approach to extract the non-parametric and parametric information of an internal clock (such as the initial phase and velocity of the internal clock) which is determined by the recent (environmental) history. We can further study how the after-effects of previous environments influence the organism’s response to the present environment. In future work, we could study how organisms switch light response modes in different external environments.

Our finding in the current study (S1 Data and S3 Fig) and the earlier report by Aschoff & Pohl (Aschoff and Pohl, 1978) suggest connections between the oscillator types and the zeitgeber strength. Weak oscillators are often observed in unicellular organisms and plants. These organisms tend to show a high sensitivity to zeitgebers. Consequently, a large entrainment range and a small slope of the POE verses period mismatch is observed. On the contrary, mammals and birds more frequently have strong oscillators with low sensitivity to zeitgebers. Thus, they have a narrow entrainment range and a steep slope of the POE verses period mismatch. An interesting observation by Aschoff & Pohl was that birds become re-entrained much faster than most mammals, although their internal clocks are both characterized as strong oscillators with similar responsivity to zeitgeber (Aschoff and Pohl, 1978). Our dCiRC model provides a possible explanation and a way to verify by fitting to available data in the future. The dCiRC model shows that the re-entrained phase is determined by both *c*_*z*_ and *k*_*e*_. With similar *c*_*z*_ values, we would expect a smaller elastic effect (*k*_*e*_) and relatively larger range of entrainment for birds.

In summary, dCiRC describes many characteristics of circadian entrainment consistent with elements of both parametric and non-parametric forms. dCiRC determines the genotype specific CiRC shapes by fitting real data. By characterizing dCiRC of 26 ecotypes, we found non-uniform light responses of these clocks. One of the major advantages of dCiRC over PRC is that dCiRC allows one to study the real-time dynamic behaviors of the clock in cycling natural environments. dCiRC embodies the concept of PRC by generating the phase shift trajectory and PRC. Conversely, we can find the genotype-specific or species-specific dCiRC curves with recorded PRCs. The significance of this work is that we can further build a “circadian fingerprint” database composed of genotype-specific or species-specific dCiRC parameters and reveal the entrainment mechanisms for various organisms. As an essential improvement, dCiRC can comprehensively study the period-phase relationship in cycling conditions. In the case when the organism entrains to environmental cues, the dCiRC model tracks the evolution of the phase relationship between the circadian internal clock and zeitgeber cycle and predicts the stable phase of entrainment. In the case that the organism is not able to entrain, dCiRC generates the phase shift trajectory to help us to investigate possible reasons causing failure of entrainment. We believe that dCiRC could allow us to study the ecology and the function of circadian clock in the natural world.

## Acknowledgements

The authors are grateful to Professor Dr. Till Roenneberg for fruitful discussions and comments on the manuscript. We thank Drs. Glass, Crosthwaite, and Larrondo for sharing Neurospora strains, and to Mr. Helmut Klausner and Ms. Angela Meckl for technical support, and to Charot Rodeget for critical reading and insightful suggestions. The work was supported by the Center of Advanced Studies of the LMU Munich and Rutgers Global International Travel Grant for generous support (to KL). The funders had no role in study design, data collection and analysis, decision to publish, or preparation of the manuscript.

## Data availability statement

All relevant experimental data are within the paper, its Supporting Information files.

The source code and data used to produce the results and analyses presented in this manuscript is available at the following public repositories,

- GitHub repository at https://github.com/AZM1994/dCiRC-Project
- Zenodo repository: <u>10.5281/zenodo.5077945</u>

## Supporting Information

**S1 Fig. Optimizing luciferase reporter in DD and LD condition. (**A) Media effect on circadian period in DD. HG1: high glucose media 1, HG2: high glucose media 2, LG: low glucose media, FGS: sorbose media. The mean value is the average of six different period algorithms in BioDare2. Error bars represent standard deviation. (B) FRQ:LUC reporter rhythm in LD. Shade areas represent the dark phase. (C) Variation Media effect in phase of entrainment in LD. (D) Media effects in amplitude. See detail description in the text.

**S2 Fig. Examples of dCiRC curves with different combinations of (*s, a*)**. is the shape factor determining the length of the flat zone around the noon. *a* is the asymmetry factor determining the ratio between the positive and negative areas. (A), (B), (C), and (D) are four illustrative examples to show how the factors (*s, a*) determine the shape of the dCiRC curves. (E), (F), (G), and (H) are four examples of the best fitting dCiRC curves for *Neurospora crassa* ecotypes. The (*s, a*) combinations for (A), (B), (C), (D), (E), (F), (G), and (H) are (0.00,1.00), (2.00,1.00), (0.50,2.00), (0.50,0.50), (0.60,0.30), (0.40,0.25), (0.00,0.57), and (0.85,0.60), respectively. The area under the curve with positive (negative) CiRC values represents the circadian system’s capacity to compress (expand) the internal period length (*τ*_*I*_). The sum of the positive and negative areas reflects the overall effect of the zeitgeber signal. A positive (negative) sum reflects the overall effects is compressing (*τ*_*I*_ (expanding (*τ*_*I*_).

**S3 Fig. Diagrams defining the total area under curve (TAUC) and light-exposed area under curve (LAUC)**. Using genotype JW238 as an example, the best fitting (*s, a*) = (0.00,0.57), A. TAUC is calculated as the total areas under the dCiRC curve (red slash shaded areas) with a sign, the area is positive from 0h to 12h, and negative from 12h to 24h. B. LAUC is calculated as the light-exposed areas under the dCiRC curve (red slash shaded areas) with a sign.

**S4 Fig. Relationship between the best fitting zeitgeber strength (*c***_***z***_ **) and the simulated phase of entrainment (Ψ**_***sim***_). The boxplot on the right shows the summary statistics for *c*_*z*_. The blue (red) area of the color bar corresponds to the small (large) values of *c*_*z*_, respectively. The scatter plot on the left shows the results of best fitting *c*_*z*_ and Ψ_*sim*_ for all 25 ecotypes. The red dotted line at *c*_*z*_ = 0.296 (0.568) represents the 25th (75th) percentile for all 25 best fitting *c*_*z*_ values, respectively. The internal clock with large *c*_*z*_ (beyond 75th percentile) is defined as the weak oscillator, the internal clock with small *c*_*z*_ (below 25th percentile) is defined as the strong oscillator.

**S1 Data. Using the dCiRC model to find best-fitting parameters of each Neurospora strain**. We performed the entrainment experiment (in DD and LD) and recorded the developmental rhythms data for 25 *N. crassa* ecotypes and DBP338. To determine the best-fitting dCiRC parameters for each strain, we fitted the simulated dCiRC phase trajectory to the processed LD phase trajectory data and solved a convex optimization problem. In the table, *τ*_*DD*_ is the free-running period measured in DD conditions, *c*_*z*_ ∈ [0,3] is the zeitgeber strength, *k*_*e*_ ∈ [0,3] is the elastic factor, *s* ∈ [0,2] is the shape factor, *a* ∈ [0,2] is the asymmetry factor.

