## Supplementary material for "A Unified Model for Entrainment by Circadian Clocks: Dynamic Circadian Integrated Response Characteristic (dCiRC)": S1 Fig

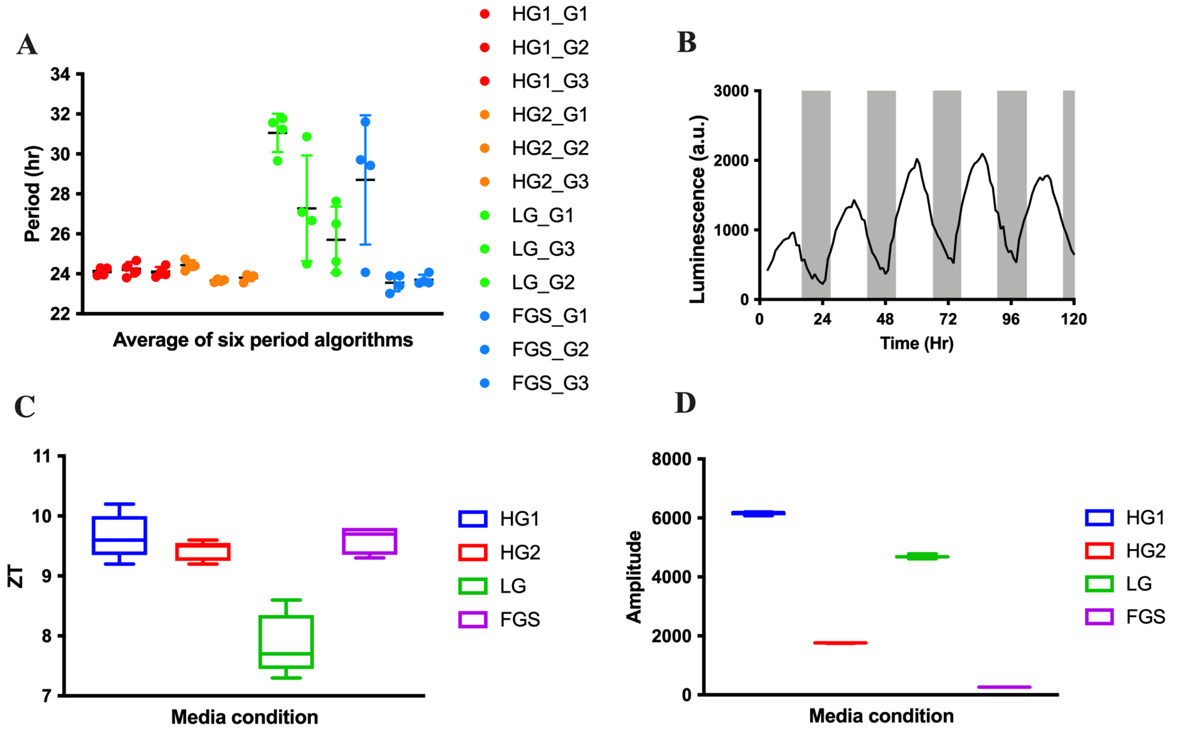


**S1 Fig. Optimizing luciferase reporter in DD and LD condition.**
**(**A) Effect of media on circadian period in DD. HG1: high glucose media 1, HG2: high glucose media 2, LG: low glucose media, FGS: sorbose media. The mean value is the average of six different period algorithms in BioDare2. Error bars represent standard deviation. (B) FRQ:LUC reporter rhythm in LD. Shaded areas represent the dark phase. (C) Effect of media on phase of entrainment in LD. (D) Effect of media on amplitude. See detail description in the text.

The translational reporter FRQ:LUC has been used for measuring the circadian rhythms in gene expression in constant dark conditions in previous studies. However, it is technically challenging to measure luciferase reporter activity under the LD condition. To optimize the assay, we performed the luciferase assay under different light conditions (DD and LD) using four culture media. We also compared six different algorithms calculating period and phase using BioDare2, biodare2.ed.ac.uk.

**Media**
We prepared media based on the 'Neurospora online protocol guide' at Fungal Genetics Stock Center; http://fgsc.net/Neurospora/NeurosporaProtocolGuide.htm
Briefly, High Glucose (HG1) media contains 2% glucose, 1X Vogel media. The salts and glucose solutions were autoclaved separately in HG1. High Glucose (HG2) media contains 2% glucose, 1X Vogel media. The salts and glucose were autoclaved together in HG2. Low Glucose (LG) media contains 0.3% glucose, 1X Vogel media. Sorbose (FGS) media contains 1% w/v L-Sorbose, 0.05% w/v D-Glucose, 0.05% w/v D-Fructose, 1x Biotin and 1X Vogel. All media are prepared as liquid media without agar.

**Plate Design**

To test the positional effect, we tested the luciferase activities at three different locations of the plate with four replicate wells of a same media.

| | 1 | 2 | 3 | 4 | 5 | 6 | 7 | 8 | 9 | 10 | 11 | 12 |
|---|---|---|---|---|---|---|---|---|---|---|---|---|
| A | HG1-1 | HG1-2 | HG1-3 | HG1-4 | LG-1 | LG-2 | LG-3 | LG-4 | FGS-9 | FGS-10 | FGS-11 | FGS-12 |
| B | HG2-1 | HG2-2 | HG2-3 | HG2-4 | | | | | | | | |
| C | HG1 | | | | HG1-5 | HG1-6 | HG1-7 | HG1-8 | HG2 | | | |
| D | HG1+Luciferin | | | | HG2-5 | HG2-6 | HG2-7 | HG2-8 | HG2+luciferin | | | |
| E | LG | | | | LG-5 | LG-6 | LG-7 | LG-8 | FGS | | | |
| F | LG+Luciferin | | | | FGS-5 | FGS-6 | FGS-7 | FGS-8 | FGS+luciferin | | | |
| G | | | | | | | | | HG2-9 | HG2-10 | HG2-11 | HG2-12 |
| H | FGS-1 | FGS-2 | FGS-3 | FGS-4 | LG-9 | LG-10 | LG-11 | LG-12 | HG1-9 | HG1-10 | HG1-11 | HG1-12 |

The four replicates of each location are grouped as group 1, e.g., HG1-G1 represents data from HG1-1, HG1-2, HG1-3, and HG1-4. Controls consisted of media with or without luciferin and with or without cells.

**Luciferase Assay**

A standard 96-well plate was used for the top counting fluorescence reader Twinkle (LB960, Berthold Tech. GmbH). 200 ml of medium was added per well containing D-luciferin (PJK GmbH #102111), for a final concentration of 10 μM from the 10 mM stock solution. Each well contained 5 x $10^5$ 7-day old conidia grown in a minimal media slant. The plate was sealed with a film that allows a gas exchange (Breath-Easy BEM-1, Diversified Biotech). The fluorescence reader was placed in a chamber with a constant temperature at 25 °C.

**Data analysis**

The raw luciferase data were uploaded into the BioDare2. To measure the variation of the period calculation by different period algorithms, we plotted the mean of the period and variation in S1A. Raw data and analysed data are available in BioDare2, Experiment# 13291. High-glucose media supported a consistent expression of luciferase activity. HG1 medium provided a more consistent luciferase reporter signal than that of HG2 after the fourth day. LG had no signal until the second day. From the third day, the signal increased significantly. However, LG provided a less consistent signal than that of HG1 or HG2. The luciferase signal in FGS medium showed a dampened signal. Almost no rhythmic signal detected from the fourth day. The luciferase signal in LG showed a significant variation in period calculation depending on the algorithm used. The luciferase signal in FGS showed a positional effect in period calculation.

To measure the luciferase activity in entrainment conditions, we tested if we could read the luciferase signal once each hour, with the plate sitting out of the machine (exposed to light or dark) between readings. The plate assay machine was placed in a chamber programmed for a 12 hr light and 12 hr dark cycling condition. We could successfully measure the rhythmic FRQ:LUC activity in LD condition ( Fig. S1B). HG and FGS media supports rhythmic luciferase reporter in LD (Fig. S1C). LG media caused an advanced phase (Fig. S1C). FGS supports robust rhythmic luciferase activity in LD, however, the amplitude of the rhythm is significantly lower than those in HG (Fig. S1D). Raw data and analyses data are available in BioDare2, Experiment# 13331.
