## Supplementary material for "A Unified Model for Entrainment by Circadian Clocks: Dynamic Circadian Integrated Response Characteristic (dCiRC)": S2 Fig

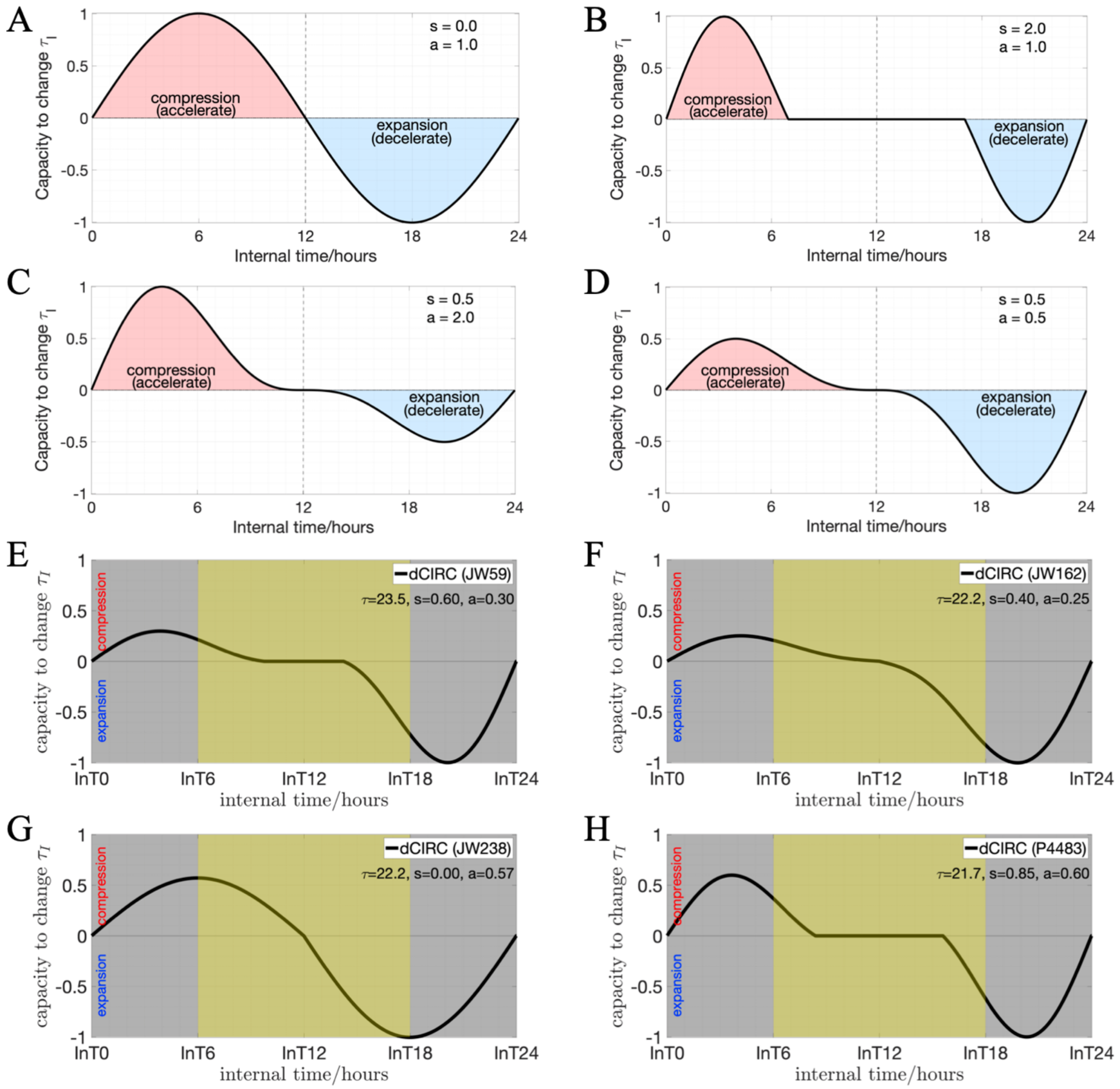


**S2 Fig. Examples of dCIRC curves with different combinations of $(s, a)$.** $s$ is the shape factor determining the length of the flat zone around the subjective noon. $a$ is the asymmetry factor determining the ratio between the positive and negative areas. (A), (B), (C), and (D) are four illustrative examples to show how the factors $(s, a)$ determine the shape of the dCIRC curves. (E), (F), (G), and (H) are four examples of the best fitting dCIRC curves for *Neurospora crassa* ecotypes. The $(s, a)$ combinations for (A), (B), (C), (D), (E), (F), (G), and (H) are $(0.00,1.00)$, $(2.00,1.00)$, $(0.50,2.00)$, $(0.50,0.50)$, $(0.60,0.30)$, $(0.40,0.25)$, $(0.00,0.57)$, and $(0.85,0.60)$, respectively. The area under the curve with positive (negative) CIRC values represents the circadian system's capacity to compress (expand) the internal period length ($\tau_I$). The sum of the positive and negative areas reflects the overall effect of the zeitgeber signal. A positive (negative) sum reflects the overall effects is compressing $\tau_I$ (expanding $\tau_I$).
