## Supplementary material for "A Unified Model for Entrainment by Circadian Clocks: Dynamic Circadian Integrated Response Characteristic (dCiRC)": S3 Fig

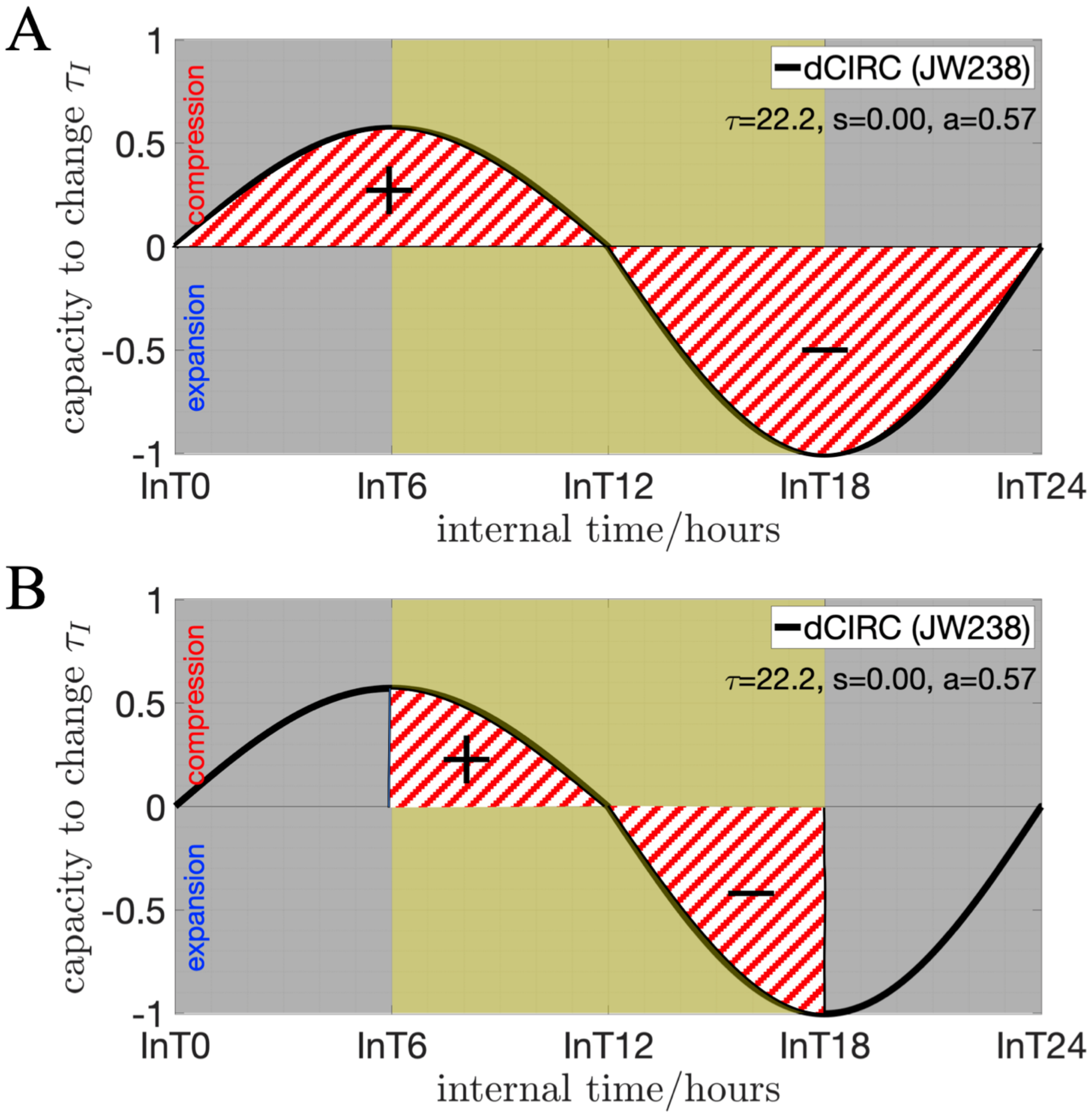


**S3 Fig.** Diagrams defining the total area under curve (TAUC) and light-exposed area under curve (LAUC). Using genotype JW238 as an example, the best-fitting $(s, a) = (0.00{,}0.57)$, A. TAUC is calculated as the total areas under the dCiRC curve (red slash shaded areas) with a sign, the area is positive from 0h to 12h, and negative from 12h to 24h. B. LAUC is calculated as the light-exposed areas under the dCiRC curve (red slash shaded areas) with a sign.
