## Supplementary material for "A Unified Model for Entrainment by Circadian Clocks: Dynamic Circadian Integrated Response Characteristic (dCiRC)": S4 Fig

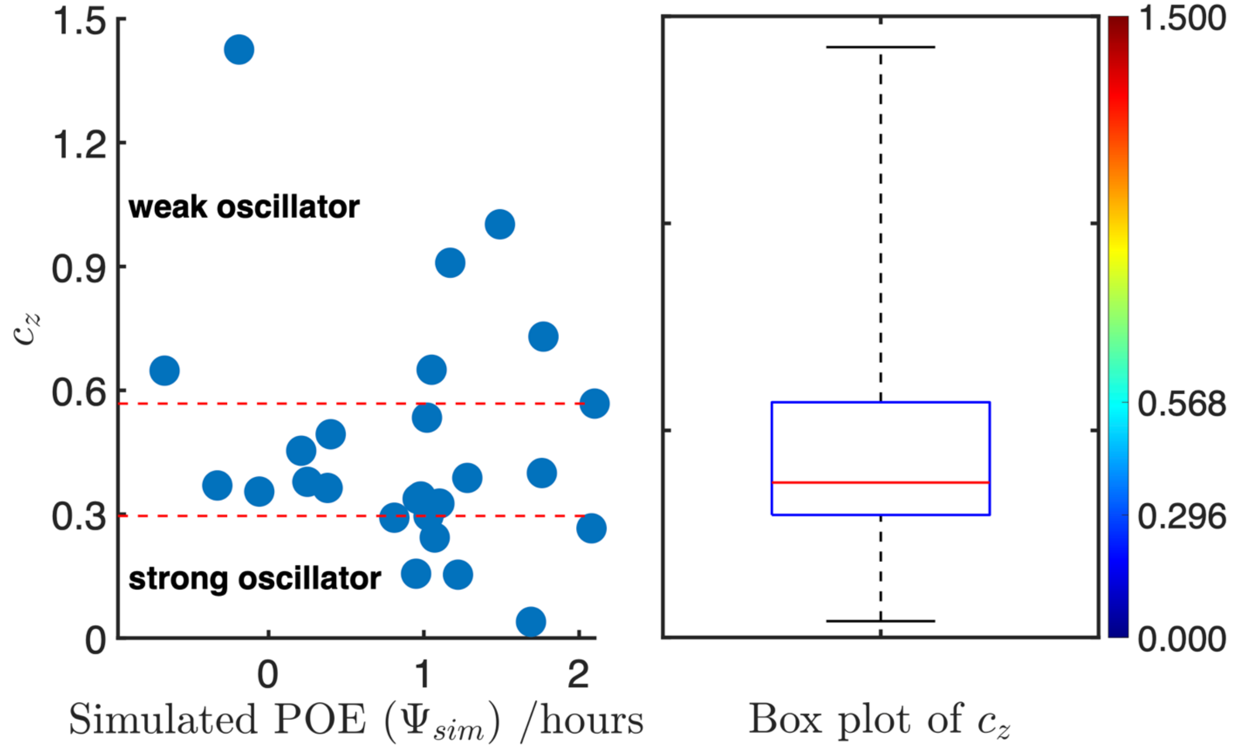


**S4 Fig. Relationship between the best fitting zeitgeber strength ($c_z$) and the simulated phase of entrainment ($\Psi_{sim}$).** The boxplot on the right shows the summary statistics for $c_z$. The blue (red) area of the color bar corresponds to the small (large) values of $c_z$, respectively. The scatter plot on the left shows the results of best fitting $c_z$ and $\Psi_{sim}$ for all 25 ecotypes. The red dotted line at $c_z = 0.296(0.568)$ represents the 25th (75th) percentile for all 25 best fitting $c_z$ values, respectively. The internal clock with large $c_z$ (beyond 75th percentile) is defined as the weak oscillator, the internal clock with small $c_z$ (below 25th percentile) is defined as the strong oscillator.
