## Supplementary material for "A Unified Model for Entrainment by Circadian Clocks: Dynamic Circadian Integrated Response Characteristic (dCiRC)": S1 Data

| | $\tau_{DD}$ | $c_z$ | $k_e$ | $s$ | $a$ | $\Psi_{sim}$ | $\Psi_{exp}$ |
|---|---|---|---|---|---|---|---|
| **JW168** | 21.0 | 0.6477 | 1.5737 | 1.40 | 0.05 | -0.97 | -0.80 |
| **JW224** | 21.0 | 0.2436 | 0.7246 | 0.00 | 0.05 | 1.07 | 1.10 |
| **JW60** | 21.0 | 0.3696 | 1.5247 | 0.05 | 0.10 | -0.33 | -0.70 |
| **JW260** | 21.2 | 0.3548 | 1.9021 | 0.00 | 0.10 | -0.06 | 0.60 |
| **D117** | 21.3 | 1.4254 | 1.7593 | 1.05 | 0.05 | -0.19 | 0.30 |
| **D119** | 21.3 | 1.0021 | 1.9621 | 0.00 | 0.05 | 1.49 | 1.00 |
| **D116** | 21.5 | 0.2958 | 0.6504 | 0.05 | 0.05 | 1.03 | 0.90 |
| **JW18** | 21.7 | 0.2658 | 0.4019 | 0.20 | 0.10 | 2.08 | 2.20 |
| **JW172** | 21.7 | 0.3373 | 0.3216 | 1.00 | 0.05 | 0.82 | 1.10 |
| **P4483** | 21.7 | 0.1559 | 0.1701 | 0.85 | 0.60 | 0.95 | 0.90 |
| **JW220** | 21.8 | 0.1536 | 0.1060 | 1.40 | 0.40 | 1.22 | 1.80 |
| **JW200** | 22.0 | 0.5679 | 0.5591 | 0.30 | 0.10 | 2.10 | 1.80 |
| **JW162** | 22.2 | 0.9090 | 1.0826 | 0.40 | 0.25 | 1.17 | 1.20 |
| **JW176** | 22.2 | 0.3997 | 1.1089 | 0.10 | 0.25 | 1.76 | 2.10 |
| **JW238** | 22.2 | 0.3784 | 1.9092 | 0.00 | 0.57 | 0.25 | 0.30 |
| **JW261** | 22.2 | 0.4937 | 1.9138 | 0.05 | 0.15 | 0.40 | 0.30 |
| **JW180** | 22.3 | 0.3879 | 1.0788 | 0.20 | 0.40 | 1.28 | 1.70 |
| **JW169** | 22.5 | 0.3429 | 0.7786 | 0.15 | 0.20 | 0.98 | 0.40 |
| **JW228** | 22.7 | 0.6499 | 1.9883 | 0.00 | 0.10 | 1.44 | 1.30 |
| **P4463** | 22.7 | 0.3258 | 0.7034 | 0.26 | 0.35 | 1.10 | 1.20 |
| **JW24** | 23.2 | 0.2915 | 0.9880 | 0.20 | 0.10 | 1.27 | 0.80 |
| **JW59** | 23.5 | 0.5339 | 0.6208 | 0.60 | 0.30 | 1.02 | 0.90 |
| **JW161** | 23.8 | 0.7301 | 1.2864 | 0.00 | 0.05 | 1.97 | 1.50 |
| **P4469** | 23.8 | 0.4537 | 0.8656 | 0.60 | 0.20 | 0.68 | 0.20 |
| **JW22** | 22.0 | 0.3636 | 1.1602 | 1.00 | 0.14 | 0.38 | 0.50 |
| **DBP338** | 30.0 | 0.0394 | 0.0056 | 1.40 | 1.85 | 1.69 | 1.67 |

**S1 Data. Using the dCIRC model to find best-fitting parameters of each Neurospora strain.** We performed the entrainment experiment (in DD and LD) and recorded the developmental rhythms data for 25 *N. crassa* ecotypes and DBP338. To determine the best-fitting dCIRC parameters for each strain, we fitted the simulated dCIRC phase trajectory to the processed LD phase trajectory data and solved a convex optimization problem. In the table, $\tau_{DD}$ is the free-running period measured in DD conditions, $c_z \in [0,3]$ is the zeitbeger strength, $k_e \in [0,3]$ is the elastic factor, $s \in [0,2]$ is the shape factor, $a \in [0,2]$ is the asymmetry factor.
